# Optogenetic tools for inducing organelle membrane rupture

**DOI:** 10.1101/2024.08.13.607738

**Authors:** Yuto Nagashima, Tomoya Eguchi, Ikuko Koyama-Honda, Noboru Mizushima

## Abstract

Disintegration of organelle membranes induces various cellular responses and has pathological consequences, including autoinflammatory diseases and neurodegeneration. Establishing methods to induce membrane rupture of organelles of interest is essential to analyze the downstream effects of membrane rupture; however, the spatiotemporal induction of rupture of specific membranes remains challenging. Here, we develop a series of optogenetic tools to induce organelle membrane rupture by using engineered Bcl-2-associated X protein (BAX), whose primary function is to form membrane pores in the outer mitochondrial membrane (OMM) during apoptosis. When BAX is forced to target mitochondria, lysosomes, or the endoplasmic reticulum (ER) by replacing its C-terminal transmembrane domain (TMD) with organelle-targeting sequences, the BAX mutants rupture their target membranes. To regulate the activity of organelle-targeted BAX, the photosensitive light-oxygen-voltage-sensing 2 (LOV2) domain is fused to the N-terminus of BAX. The resulting LOV2–BAX fusion protein exhibits blue light–dependent membrane-rupture activity on various organelles, including mitochondria, the ER, and lysosomes. Thus, LOV2–BAX enables spatiotemporal induction of membrane rupture across a broad range of organelles, expanding research opportunities on the consequences of organelle membrane disruption.

## Introduction

Eukaryotic cells are compartmentalized by organelle membranes, and the disintegration of these membranes triggers numerous biological responses and is associated with various diseases. Excessive release of mitochondrial intermembrane proteins, such as cytochrome *c*, from ruptured mitochondria activates apoptosis signaling through the activation of caspases (1, 2). In cases of minor membrane permeabilization that are insufficient to cause cell death, activated caspases may induce DNA damage and promote oncogenesis (3). Mitochondrial membrane rupture also causes leakage of mitochondrial DNA (mtDNA) (4). Cytoplasmic mtDNA is sensed by cyclic GMP-AMP synthase, activates stimulator of interferon genes (STING), and subsequently initiates inflammatory signaling. These downstream processes are involved in cancers, autoimmune diseases, and neurodegenerative disorders (5, 6). Lysosomal membrane rupture also leads to cytotoxicity via leakage of hydrolysis enzymes into the cytosol (7). Because of the undesirable consequences of membrane disintegration, cells have protective mechanisms, including autophagy-mediated clearance(8-10), and the endosomal sorting complex required for transport (ESCRT) or annexin-dependent repair of ruptured organelles (11, 12). Organelle membrane rupture can also occur under normal physiological conditions. For instance, in lens fiber cells, organelle membranes are first slightly damaged by unknown mechanisms during development and can then be fully degraded by phospholipase A/acyltransferase (PLAAT) family phospholipases, leading to lens transparency (13).

To analyze the molecular responses to organelle membrane rupture and their eventual pathophysiological consequences, methods to induce rupture of specific organelle membranes are essential. However, at present spatiotemporal induction of membrane rupture remains a challenge. Although chemical compounds such as B cell lymphoma-2 (Bcl-2) inhibitors and L-leucyl-L-leucine methyl ester (LLOMe) are widely used to rupture the mitochondrial and lysosomal membranes, respectively (14, 15), they cannot induce membrane rupture in specific cells within a tissue or in specific organelles of a single cell. In addition, chemicals that rupture membranes of the endoplasmic reticulum (ER), peroxisomes, or the Golgi apparatus are not yet available. Several studies have reported optogenetic tools to damage organelles (16-18). Although these tools can spatiotemporally induce membrane rupture, their targets are limited to mitochondria alone (16, 17) and both mitochondria and peroxisomes (18).

Bcl-2-associated X protein (BAX) is a Bcl-2 family protein that mediates permeabilization of the outer mitochondrial membrane (OMM) during apoptosis (19). BAX is classified as a tail-anchored protein that possesses a single transmembrane domain (TMD) in its C-terminal region, followed by a short C-terminal element. In its inactive state, the C-terminal region of BAX is concealed within the N-terminal soluble domain, resulting in BAX being retained in the cytosol (20). Upon receiving apoptotic stimuli, the C-terminal region of BAX becomes exposed, and BAX is then inserted into the OMM, where BAX, along with another proapoptotic protein, Bcl-2 homologous antagonist/killer (BAK), undergoes oligomerization to form a membrane pore (21). It is reported that artificial targeting of BAX or BAK to mitochondria or ER results in leakage of their luminal contents (22).

In this study, by engineering BAX, we developed a series of optogenetic tools wherein photoactivatable BAX is specifically targeted to the membranes of distinct organelles. Using these tools, we were able to rupture the mitochondrial, ER, and lysosomal membranes in a light-dependent manner.

## Results

### Organelle-targeted BAX ruptures the mitochondrial, lysosomal, and ER membranes

To establish inducible organelle rupture systems, we first prepared reporters to detect membrane rupture of various organelles. As a marker for mitochondrial membrane rupture, HaloTag (hereafter referred to as Halo)–PLAAT3(C113S), a catalytically inactive mutant of mouse PLAAT3, was used (Fig. 1A). PLAAT3 is a lipase that translocates from the cytosol to damaged organelles such as mitochondria (however, its precise mechanism is not yet known) (13). As a marker for lysosomal membrane rupture, we used galectin-3 (Gal3) (Fig. 1A), a cytosolic protein that accumulates in ruptured lysosomes by binding to glycosylated proteins along the inner surface of the lysosomal membrane and in the lysosomal lumen (23). To visualize the ruptured ER, we utilized the FK506-binding protein (FKBP) and FKBP-rapamycin binding protein (FRB) tags, which form a heterodimer in the presence of A/C heterodimerizer (hereafter simply referred to as heterodimerizer), a derivative of rapamycin (24). FRB was expressed in the ER lumen by fusing it to the C-terminus of SEC61B, while FKBP–Halo was expressed in the cytosol. Cytosolic FKBP bound to FRB and accumulated in the ER lumen, in a manner dependent on both the presence of the heterodimerizer and the rupture of ER membranes (Fig. 1A).

**Figure 1.**
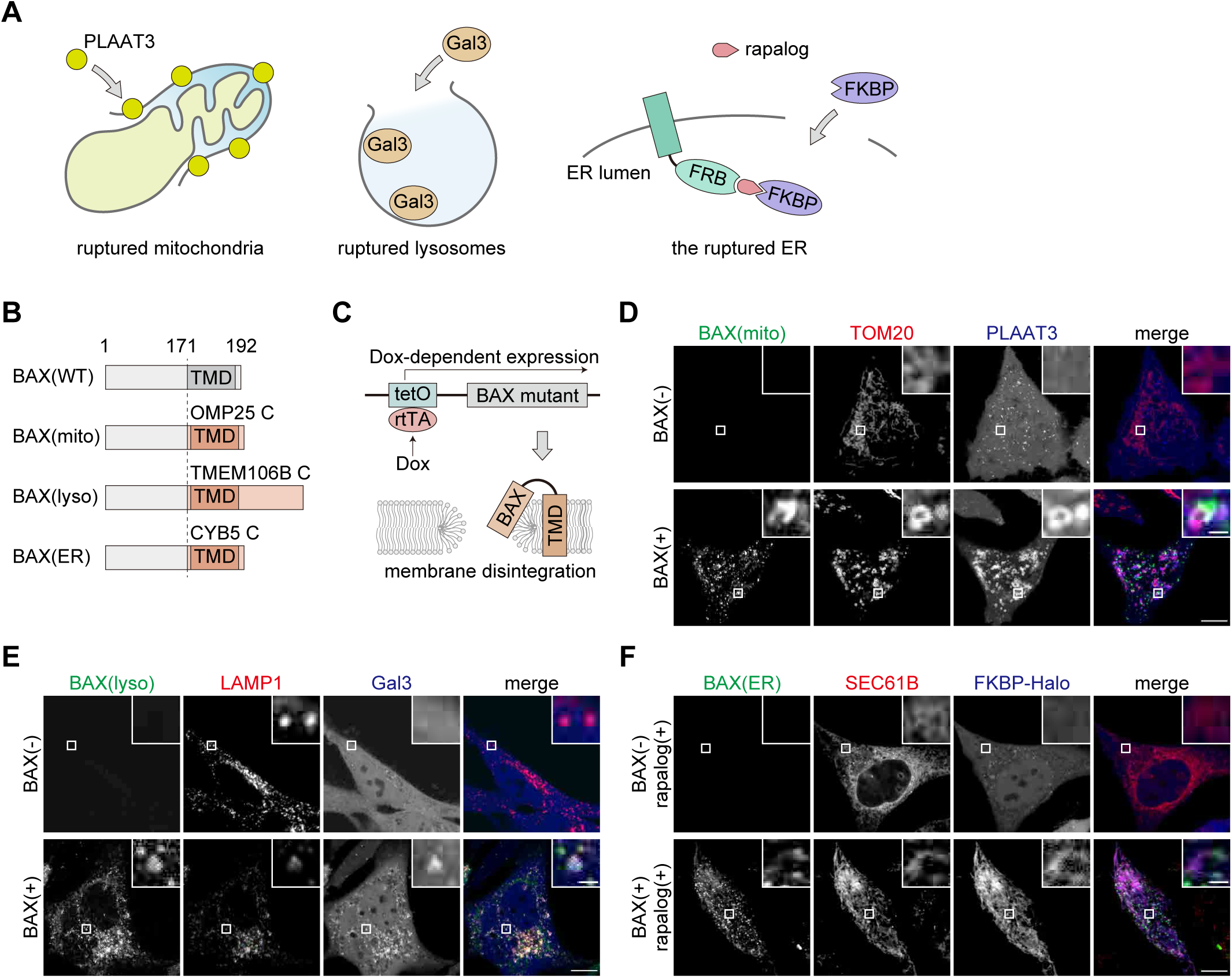
Forced targeting of BAX induces organelle membrane rupture. **A,** Reporters used to detect organelle membrane rupture. Each reporter translocates from the cytosol to its target organelles upon membrane rupture. **B,** Construction of BAX mutants. The C-terminal region of BAX was replaced with that of OMP25, TMEM106B, or CYB5. **C,** Regulation of BAX expression using a Tet-On system. Reversed tetracycline transactivator (rtTA) binds to tetracycline operator elements (tetO) and induces expression of the BAX mutants in a doxycycline-dependent manner. The BAX mutants localize to target organelles and rupture the membranes. **D,** HeLa cells expressing TOM20–mRFP and Halo–PLAAT3(C113S) without (upper panels) or with (lower panels) Tet-ON GFP–BAX(mito) were treated with doxycycline for 1 day in the presence of Q-VD-Oph. The HaloTag ligand conjugated with SaraFluor 650T was added 10 min before observation. Scale bars, 10 μm (main panels), 1 μm (inset panels). **E,** HeLa cells expressing LAMP1–mRFP and Halo–Gal3 without (upper panels) or with (lower panels) Tet-ON GFP–BAX(lyso) were treated with doxycycline for 2 days in the presence of Q-VD-Oph. HaloTag SaraFluor 650T ligand was added 10 min before observation. Scale bars, 10 μm (main panels), 1 μm (inset panels). **F,** HeLa cells expressing SEC61B–mCherry–FRB and FKBP–Halo without (upper panels) or with (lower panels) Tet-ON GFP–BAX(lyso) were treated with doxycycline for 1 day in the presence of Q-VD-Oph. The cells were treated with heterodimerizer and HaloTag SaraFluor 650T ligand for 10 min. Scale bars, 10 μm (main panels), 1 μm (inset panels).

Then, we tested whether BAX induced rupture of organelle membranes when it was forced to target mitochondria, lysosomes, and the ER by replacing its transmembrane domain (TMD) in its C-terminal region with that of respective organelle-specific proteins (Fig. 1B). Because organelle membrane rupture may cause cell death, these BAX mutants were inducibly expressed in HeLa cells in a doxycycline-dependent manner (Fig. 1C), and cells were cultured in the presence of quinoline-Val-Asp-difluorophenoxymethylketone (Q-VD-Oph), a caspase inhibitor, to block the cell death signaling pathway.

To localize BAX to mitochondria, we replaced the C-terminal region of BAX(171–192) with that of the mitochondrial tail-anchored protein OMP25(109–145), which is referred to as BAX(mito). When EGFP–BAX(mito) was not expressed, Halo–PLAAT3(C113S) was mostly diffused throughout the cytoplasm and did not colocalize with mitochondria (though some small punctate structures were observed that likely indicated peroxisomes, as previously reported (25)) (Fig. 1D). After induction of EGFP–BAX(mito) expression by doxycycline, EGFP–BAX(mito) was successfully recruited to TOM20-positive OMM. Subsequently, Halo–PLAAT3(C113S) accumulated on these mitochondria, suggesting that the mitochondrial membranes were ruptured (Fig. 1D).

To localize BAX to lysosomes, we initially utilized the TMDs of VAMP7 and VAMP8, lysosomal tail-anchored SNARE proteins; however resultant constructs failed to localize to lysosomes. Then, we constructed BAX(lyso), in which the TMD of BAX was replaced with the C-terminus (amino acids 90–274) of TMEM106B, a single-pass membrane protein on lysosomes, whose N-terminus faces into the cytosol (26). Following doxycycline treatment, EGFP–BAX(lyso) was recruited to the LAMP1-positive lysosomes, and Halo–Gal3 was translocated from the cytosol to the lysosomes (Fig. 1E), suggesting that the lysosomal membranes were indeed ruptured.

We also generated BAX(ER) by replacing the C-terminus of BAX with the C-terminus (amino acids 95–134) of CYB5, an ER transmembrane protein, to localize BAX to the ER. Upon treatment with doxycycline, EGFP–BAX(ER) localized to the ER (Fig. 1F). When EGFP–BAX(ER) was not expressed, FKBP–Halo was diffusely distributed throughout the cytosol, even in the presence of the heterodimerizer. After cells expressed EGFP–BAX(ER) and were treated with the heterodimerizer, FKBP– Halo accumulated on the ER membranes (Fig. 1F), indicating that EGFP–BAX(ER) induced ER membrane rupture.

These data indicate that forced translocation of BAX to organelle membranes induces membrane rupture regardless of organelle type.

### Tethering BAX to organelle membranes induces mitochondria-specific membrane rupture

Although the doxycycline-based system successfully ruptured organelle membranes inductively, the transcription-dependent system was not very rapid. To overcome this limitation, we sought to introduce an optogenetic system. Previous studies have demonstrated that tethering of cytoplasmic BAX to the mitochondrial membranes using optogenetic tags can be used to induce mitochondrial membrane rupture (16, 17). In this system, photoactivable dimerization modules of *Arabidopsis* CIB and CRY2 are fused to TOM20 and BAX (with mutation S184E to prevent background targeting to mitochondria), respectively. Photo-induced dimerization of CRY2 and CIB tags results in the tethering of BAX to the OMM and induces OMM rupture. To test whether a similar method can be applied to the inducible rupture of other organelles, we modified the BAX translocation system using the FKBP and FRB tags (Fig. 2A).

**Figure 2.**
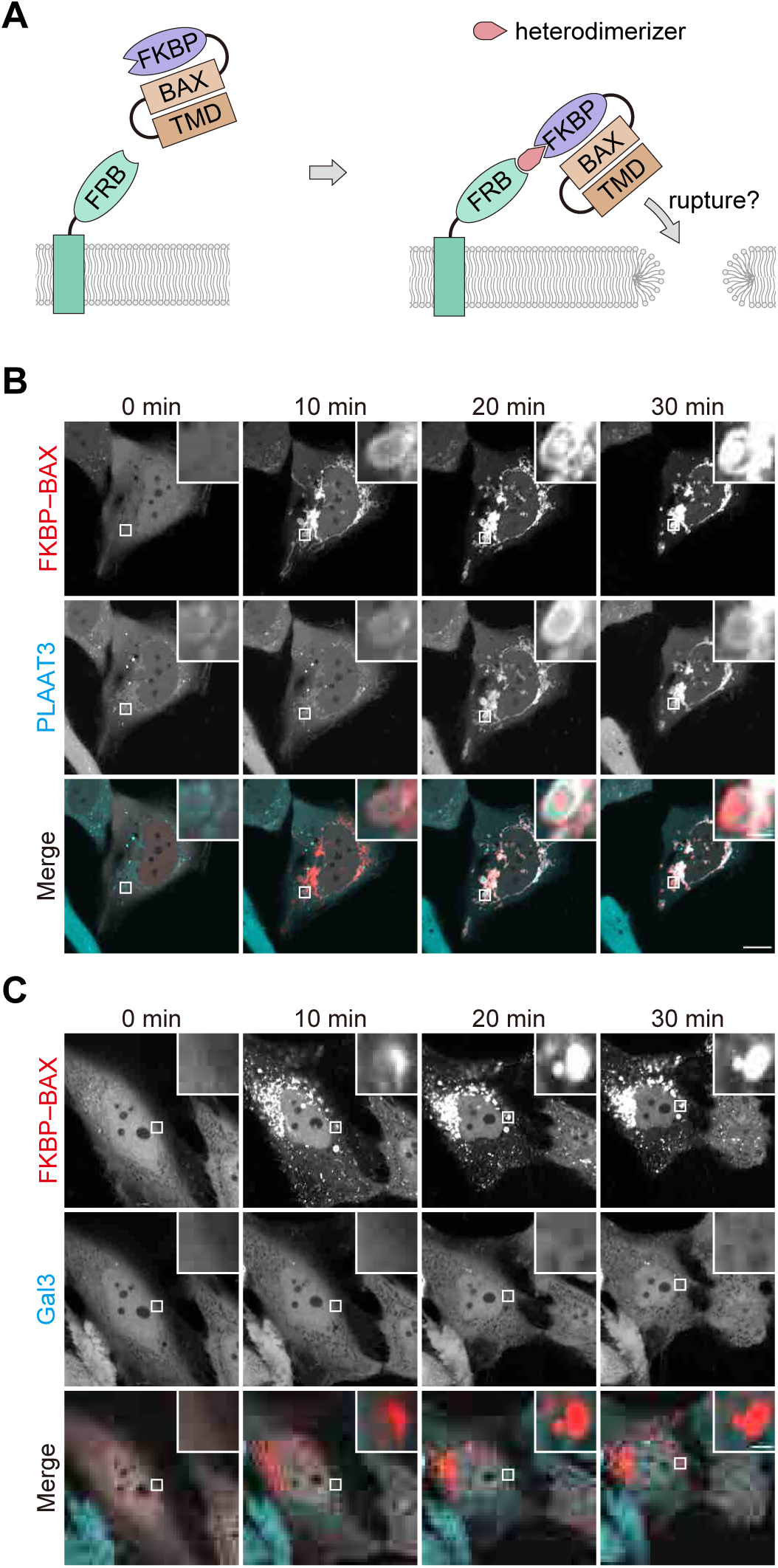
Membrane tethering of BAX causes rupture of the outer mitochondrial membrane but not the lysosomal membrane. **A,** A schematic representation of the ectopic tethering of BAX mutants to organelle membranes via the FKBP and FRB tags. FKBP–mCherry–BAX(S184E) in the cytosol and FRB on organelle membranes form a heterodimer in the presence of heterodimerizer, bringing BAX into close proximity to the organelle membranes. **B,** HeLa cells expressing FKBP–mCherry–BAX(S184E), TOM20(1-69)–FRB, and Halo–PLAAT3(C113S) were treated with heterodimerizer and Q-VD-Oph, which were added at 0 min. HaloTag SaraFluor 650T ligand was added 10 min before observation. Scale bars, 10 μm (main panels), 1 μm (inset panels). **C,** HeLa cells expressing FKBP–mCherry–BAX(S184E), LAMP1–FRB, and Halo–Gal3 were treated with heterodimerizer and Q-VD-Oph, which were added at 0 min. HaloTag SaraFluor 650T ligand was added 10 min before observation. Scale bars, 10 μm (main panels), 1 μm (inset panels).

As expected, FKBP–mCherry–BAX(S184E) was diffusely distributed throughout the cytoplasm in the absence of the heterodimerizer, whereas FKBP– mCherry–BAX(S184E) immediately accumulated on mitochondria upon heterodimerizer treatment in HeLa cells expressing TOM20–FRB (Fig. 2B). Halo– PLAAT3(C113S) also accumulated on mitochondria, indicating that anchoring of BAX induced mitochondrial rupture (Fig. 2B). However, when FKBP–mCherry– BAX(S184E) was recruited to lysosomes via LAMP1–FRB, no Halo–Gal3 accumulation in the lysosomes was observed (Fig. 2C). These data suggest that tethering of BAX is sufficient to induce membrane rupture only in mitochondria and not in lysosomes, prompting us to develop another method to induce membrane rupture.

### Mitochondria-targeted LOV2–BAX induces blue light–dependent OMM rupture

When BAX was inserted into membranes by modifying its C-terminal TMD, BAX induced membrane rupture of various organelles (Fig. 1). Therefore, we attempted to control the activity of the “membrane-inserted” BAX mutants rather than controlling BAX translocation. To this end, we used the light-oxygen-voltage-sensing 2 (LOV2) domain, a photosensor domain from phototropin1 (Fig. 3A). The LOV2 domain is composed of the N-terminal Period-ARNT-Singleminded (PAS) domain and the C-terminal Jα-helix, which tightly bind to each other and can inhibit the function of downstream fused proteins through steric hindrance (27). Upon blue light stimulation, the Jα-helix is released from the PAS domain and then unfolded (28). Because of this photo-dependent dynamic conformational change, the LOV2 domain is widely used as a photo-switch to regulate the function of proteins by fusing to their N-termini (29). By fusing LOV2(N538E), a mutant with reduced background activation (30), to the N-terminus of BAX(mito), we constructed LOV2(N538E)–BAX(3-171)–OMP25, referred to as LOV2–BAX(mito). EGFP–LOV2–BAX(mito) localized onto mitochondria irrespective of blue light irradiation (Fig. 3B). In contrast to EGFP–BAX(mito), EGFP– LOV2–BAX(mito) caused only minor accumulation of Halo–PLAAT3(C113S) on mitochondria without blue light irradiation (Fig. 3B, 3D), indicating that tagging with the LOV2 domain suppresses membrane damage activity of BAX. After blue light irradiation for 60 min and subsequent incubation for 30 min, Halo–PLAAT3(C113S) accumulated on the mitochondria (Fig. 3B, 3D). In these cells, EGFP–LOV2– BAX(mito) formed bright puncta on the OMM, probably because EGFP–LOV2– BAX(mito) accumulated at membrane pores (see Discussion). However, blue light irradiation did not induce translocation of Halo–PLAAT3(C113S) in cells expressing EGFP–LOV2–OMP25 without the BAX cytoplasmic domain, indicating that mitochondrial rupture is not caused by phototoxicity (Fig. 3C, 3D). To confirm mitochondrial rupture, we used previously reported probes based on chemically dimerizable FKBP/FRB domains with slight modifications (3). A Halo-fused FKBP domain and a mitochondria-targeted FRB domain (fused with the mitochondrial anchoring sequence of apoptosis-inducing factor [AIF]) were expressed in HeLa cells (Fig. 3E). As expected, FKBP–Halo was recruited from the cytosol to mitochondria in a manner dependent on blue light irradiation and the heterodimerizer (Fig. 3F), supporting the inference that EGFP–LOV2–BAX(mito) induces photo-dependent OMM rupture. The OMM rupture by EGFP–LOV2–BAX(mito) was further confirmed by correlative light and electron microscopy. Without blue light irradiation, mitochondria showed oval shapes with clear cristae structures (Fig. 4). After blue light irradiation, Halo– PLAAT3(C113S) accumulated on mitochondria, which were swollen with OMM rupture and cristae structure loss (Fig. 4). These data indicate that LOV2–BAX enables induction of mitochondrial membrane rupture in a blue light–dependent manner.

**Figure 3.**
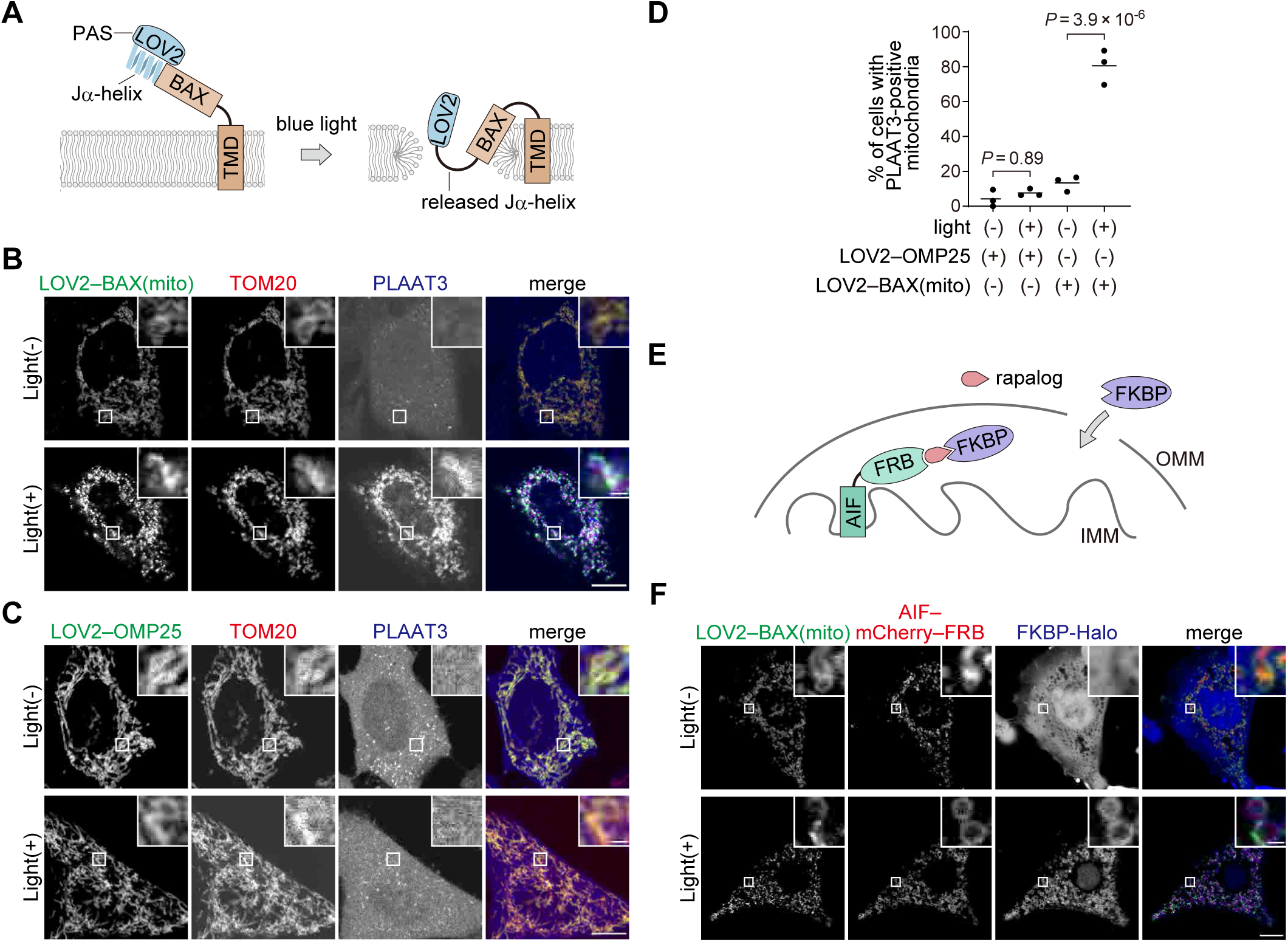
The LOV2 domain enables photo-dependent regulation of the pore-forming activity of BAX on mitochondria. **A,** The strategy used to regulate the pore-forming activity of BAX in a photo-illumination-dependent manner. LOV2 attachment inhibits the pore-forming activity of BAX. The Jα-helix of LOV2, connected to the N-terminus of BAX, is released from the PAS domain and unfolded upon blue-light irradiation. This conformational change activates the pore-forming activity of BAX. **B,** GFP–LOV2–BAX(mito), TOM20–mRFP, and Halo–PLAAT3(C113S) were expressed in HeLa cells. Cells were treated with Q-VD-Oph and HaloTag SaraFluor 650T ligand, irradiated with blue light for 60 min, and subsequently incubated for 30 min before fixation. Scale bars, 10 μm (main panels), 1 μm (inset panels). **C,** GFP–LOV2–OMP25, TOM20–mRFP, and Halo–PLAAT3(C113S) were expressed in HeLa cells. Cells were treated as in **(B)**. Scale bars, 10 μm (main panels), 1 μm (inset panels). **D,** The percentage of cells showing accumulation of Halo–PLAAT3(C113S) on the outer mitochondrial membrane. Horizontal lines indicate the group means, and each dot indicates the data point from one of three independent experiments. Differences were statistically analyzed using one-way ANOVA with Tukey’s test. At least 70 cells were observed in each experiment. **E,** A schematic illustration of the detection of mitochondrial rupture using AIFM1(1-90)–mCherry–FRB and FKBP–Halo reporters. Dependent on mitochondrial rupture and the presence of the heterodimerizer, cytosolic FKBP–Halo binds to the FRB tag located in the intermembrane space of mitochondria, resulting in the accumulation of FKBP– Halo in mitochondria. **F,** GFP–LOV2–BAX(mito), AIFM1(1-90)–mCherry–FRB, and FKBP–Halo were expressed in HeLa cells. Cells were treated with heterodimerizer, irradiated with blue light for 60 min, and subsequently incubated for 30 min before fixation. Scale bars, 10 μm (main panels), 1 μm (inset panels).

**Figure 4.**
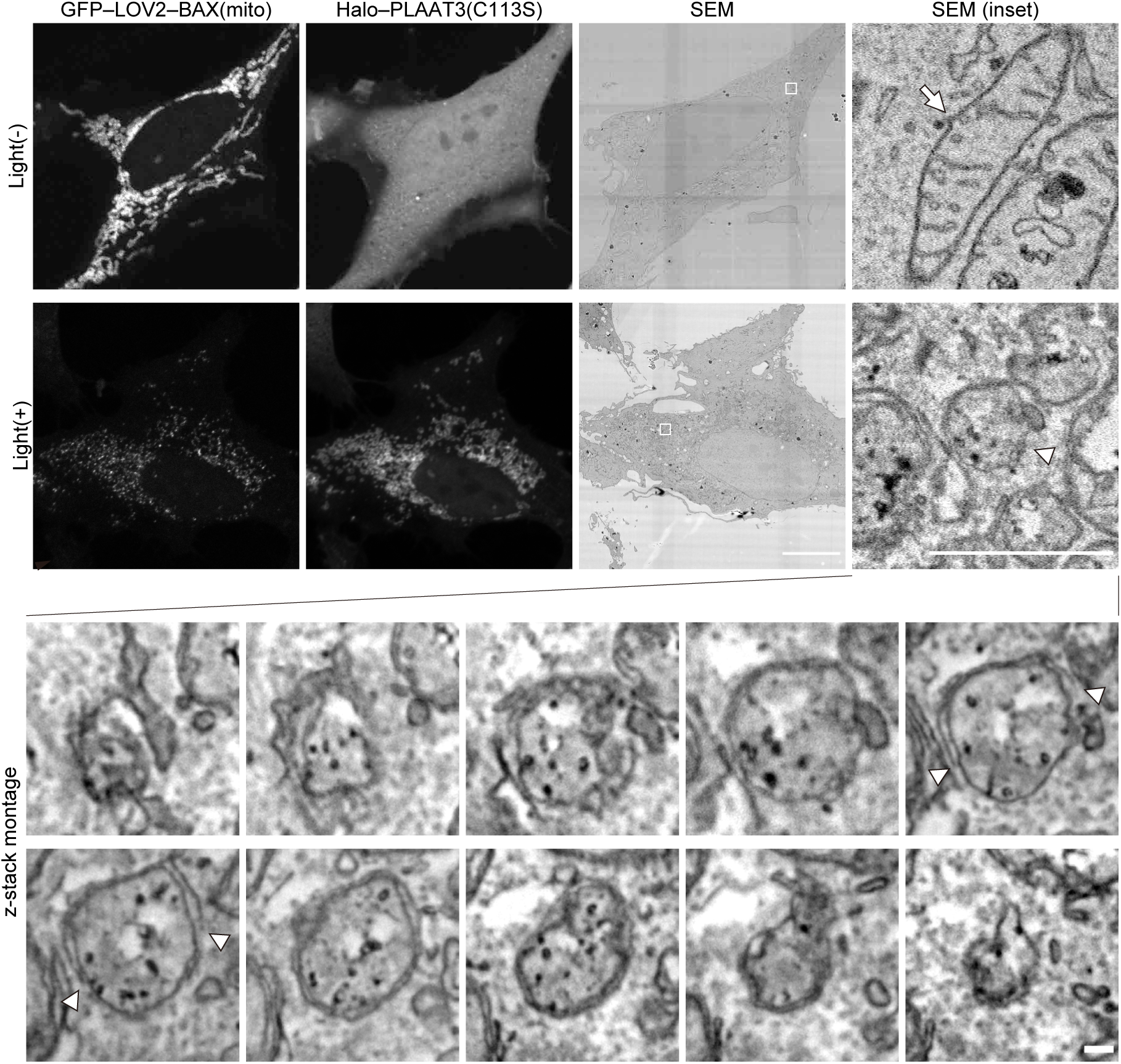
Electron micrographs of mitochondria ruptured by GFP–LOV2– BAX(mito) GFP–LOV2–BAX(mito) and Halo–PLAAT3(C113S) were expressed in HeLa cells. Cells were treated with Q-VD-Oph and HaloTag SaraFluor 650T ligand, irradiated with blue light for 60 min, subsequently incubated for 30 min before fixation, and analyzed by correlative light and electron microscopy. An arrow indicates a normal mitochondrion, and a arrowhead indicates a membrane-ruptured mitochondrion (upper panels). Serial sections (25-nm thickness) of a membrane-ruptured mitochondrion are shown (bottom panels). Arrowheads in the serial sections indicate the edge of the outer mitochondrial membrane (OMM) of the ruptured mitochondrion. SEM, scanning electron micrograph. Scale bars, 10 μm (main panels), 1 μm (enlarged panels), 100 nm (serial sections).

### LOV2–BAX induces rupture of lysosomal and ER membranes

We tested whether LOV2–BAX induced membrane rupture in organelles other than mitochondria. To induce lysosomal rupture, we first tested EGFP–LOV2(N538E)– BAX(3-171)–TMEM106B(90-274); however, this construct caused high background lysosomal damage independent of blue light. Therefore, to further reduce background activation, we introduced the additional mutation G528A into LOV2 (30) and generated EGFP–LOV2(G528A, N538E)–BAX(3–171)–TMEM106B(90–274), referred to as EGFP–LOV2–BAX(lyso). Before blue light irradiation, cells expressing EGFP–LOV2– BAX(lyso) showed only a few Gal3-positive lysosomes (Fig. 5A). The number of Gal3-positive puncta increased after photostimulation (Fig. 5A, 5C), indicating that EGFP– LOV2–BAX(lyso) triggers lysosomal rupture that is dependent on blue light. Photo-dependent lysosomal rupture was not observed in cells expressing LOV2(lyso), which lacks the BAX cytoplasmic domain (Fig. 5B, 5C).

**Figure 5.**
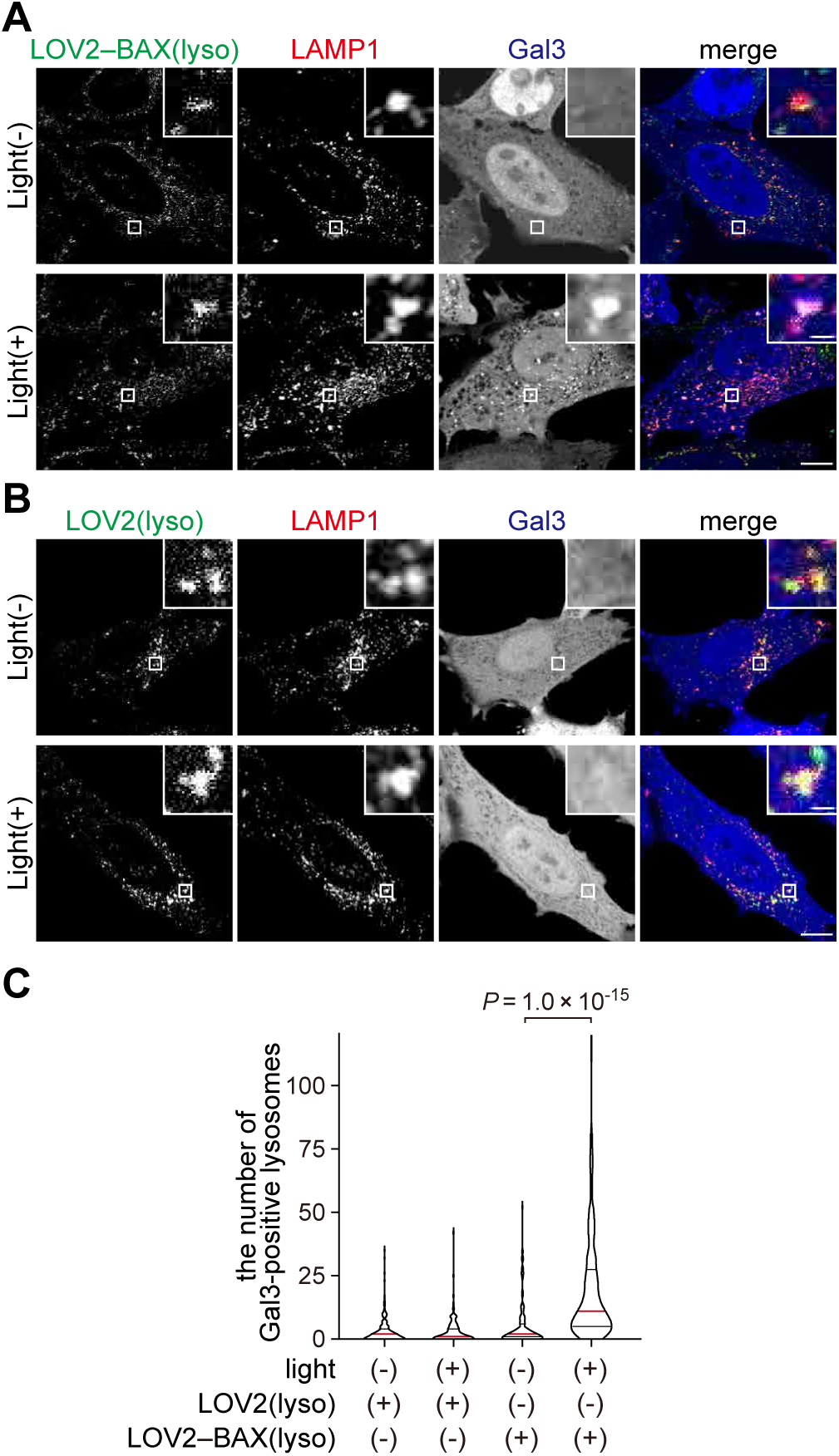
Photo-regulated lysosome rupture by LOV2–BAX. **A, B,** HeLa cells expressing LAMP1–mRFP, Halo–Gal3, and GFP–LOV2–BAX(lyso) **(A)** or GFP–LOV2–TMEM106B **(B)** were treated with Q-VD-Oph and HaloTag SaraFluor 650T ligand, irradiated with blue light for 60 min, and subsequently incubated for 30 min before fixation. Scale bars, 10 μm (main panels), 1 μm (inset panels). **C,** Quantification of the number of Halo–Gal3 puncta per cell in **(A)** and **(B)**. Differences were statistically analyzed by one-way ANOVA with Tukey’s test. The red lines represent the median, and the black lines represent the quartiles. At least 150 cells were analyzed in each experiment. **D,** Construction of BAX mutants in which the C-terminal region was replaced with that of VAMP7. The LOV2 domain was added to the N-terminus of BAX. **E,** GFP–BAX–VAMP7(171-220) was expressed in a doxycycline-dependent manner in HeLa cells constitutively expressing LAMP1–mRFP and Halo–Gal3. Scale bars, 10 μm (main panels), 1 μm (inset panels). **F,** GFP–LOV2–BAX–VAMP7(171-220), LAMP1–mRFP, and Halo–Gal3 were expressed in HeLa cells. Cells were irradiated with blue light for 60 min and subsequently incubated for 30 min before fixation. Scale bars, 10 μm (main panels), 1 μm (inset panels).

To induce photo-dependent ER rupture, we constructed EGFP– LOV2(N538E)–BAX(3–171)–CYB5(95–134), referred to as EGFP–LOV2–BAX(ER). EGFP–LOV2–BAX(ER) was expressed in HeLa cells, and ER rupture was then assessed by using the SEC61B–mCherry–FRB and FKBP–Halo reporters. EGFP– LOV2–BAX(ER) colocalized with SEC61B–mCherry–FRB, confirming ER localization of LOV2–BAX(ER) (Fig. 6A). In the presence of heterodimerizer, FKBP– Halo accumulated in the ER in a manner dependent on both LOV2–BAX(ER) and blue light irradiation (Fig. 6A–6C). Thus, these data suggest that LOV2–BAX enables blue light–dependent rupture of the membranes of mitochondria, lysosomes, and the ER.

**Figure 6.**
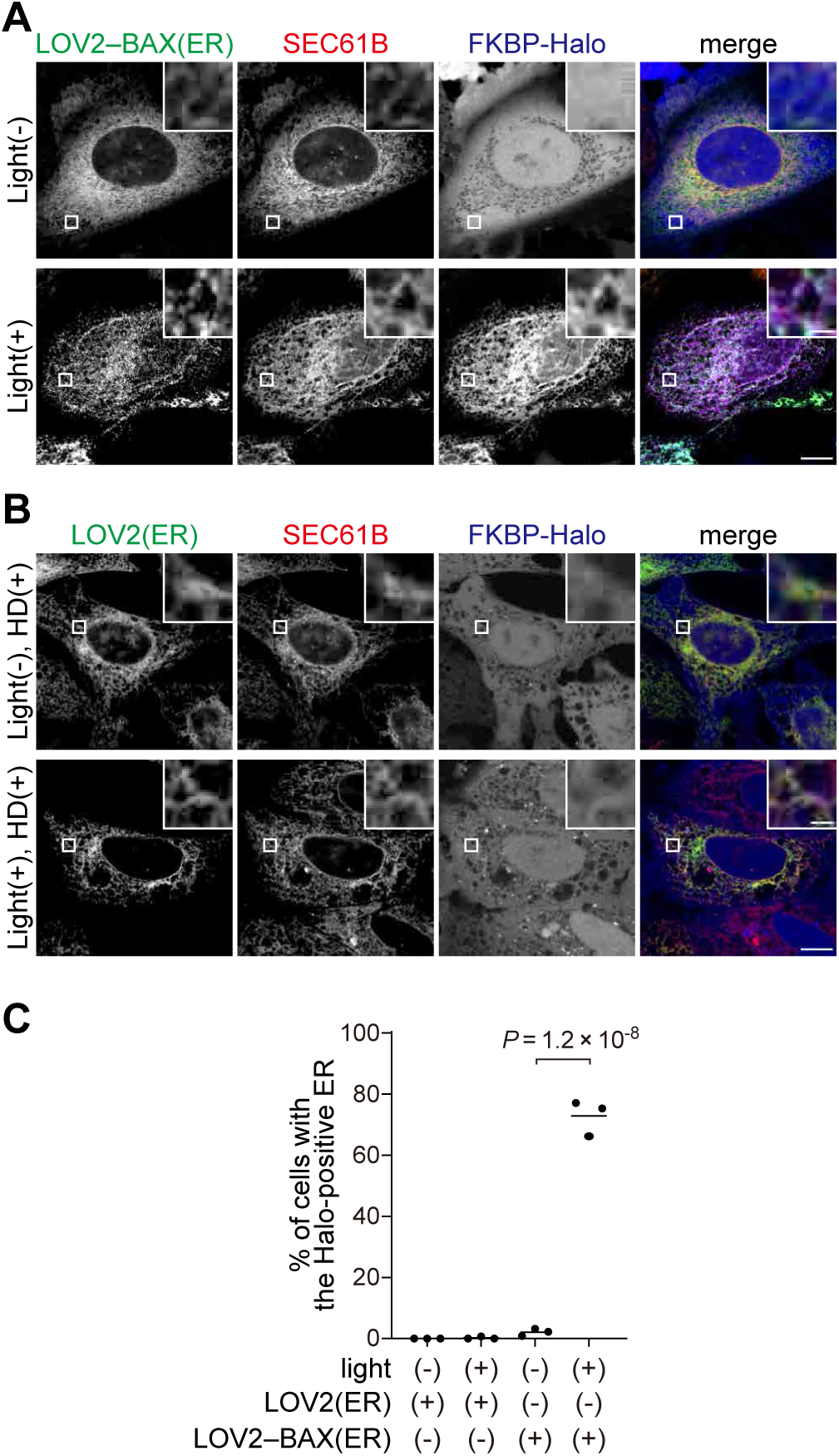
Photo-regulated ER rupture by LOV2–BAX. **A, B,** HeLa cells expressing SEC61B–mCherry–FRB, FKBP–Halo, and either GFP– LOV2–BAX(ER) **(A)** or GFP–LOV2–CYB5 **(B)** were treated with heterodimerizer, Q-VD-Oph, and HaloTag SaraFluor 650T ligand, irradiated with blue light for 60 min, and subsequently incubated for 30 min before fixation. Scale bars, 10 μm (main panels), 1 μm (inset panels). **C,** The percentage of cells showing accumulation of FKBP–Halo on the ER membrane in **(A)** and **(B)**. Horizontal lines indicate the means, and each dot indicates the data point from one of three independent experiments. Differences were statistically analyzed using one-way ANOVA with Tukey’s test. At least 100 cells were analyzed in each experiment.

## Discussion

In this study, we developed and validated a set of optogenetic tools to induce membrane rupture of various organelles. Membrane-inserted BAX can be used to induce membrane rupture regardless of the type of organelle. In conjunction with photosensitive LOV2, the membrane-rupture activity of BAX is successfully regulated in a blue light–dependent manner. This new technique enables spatiotemporal induction of membrane ruptures in specific organelles, including mitochondria, lysosomes, and the ER.

Previous studies have reported that tethering of cytoplasmic BAX to mitochondria via the optogenetic cryptochrome 2 (CRY2) and cryptochrome-interacting basic-helix-loop-helix 1 (CIB1) tags can be used to induce the rupture of the OMM (16, 17). Similar results were obtained using the FKBP and FRB tags in the present study (Fig. 2B). However, this “tethering strategy” failed to rupture the lysosomal membrane (Fig. 2C). This might be because simple tethering of BAX TMD to lysosomes is insufficient for its insertion into the lysosomal membrane, which is not the original destination of BAX. Therefore, we modified its TMD to change the localization of “membrane-inserted” BAX, enabling the regulation of pore-forming activity by using LOV2. This strategy has expanded the types of target organelles that can be ruptured by BAX. It remains to be determined whether LOV2–BAX can induce membrane rupture in other organelles, such as peroxisomes and the Golgi apparatus.

Although the regulatory mechanism of BAX by LOV2 was not analyzed in this study, LOV2 may inhibit the oligomerization of BAX, which is a key step in forming membrane pores. Oligomerized BAX and BAK accumulate around the edges of pores and are observed as puncta on the mitochondrial membranes (31). Supporting our hypothesis, the localization of EGFP–LOV2–BAX changed from a diffuse pattern to bright spots on organelle membranes upon blue light irradiation (Fig. 3B). Further research, including structural analysis, is required to reveal the precise mechanism by which LOV2 regulates BAX.

An inherent limitation of optogenetic tools is that light irradiation can cause photodamage. In this study, cells were exposed to blue light for 60 min. In this condition, we confirmed that BAX-independent membrane rupture was not observed, but cells may have been mildly damaged. To reduce phototoxicity, further optimization, including shortening of the necessary irradiation duration, is required. The wild-type LOV2 is inactivated shortly after the cessation of irradiation (about 60 s) (32, 33); therefore, continuous irradiation is required to maintain BAX activity. In addition, LOV2 mutants possessing prolonged photocycles may be useful to shorten the irradiation duration and reduce unintended phototoxicity.

Unlike chemicals such as LLOMe, optogenetic LOV2–BAX enables spatial regulation of organelle membrane rupture. The induction of membrane rupture at the single organelle level and the analysis of the cellular responses are future challenges. Spatial regulation will also benefit in vivo analysis. By regulating both the irradiation area and the expression pattern of LOV2–BAX, our tools can induce organelle membrane rupture in specific tissues. Investigating in vivo responses to organelle membrane rupture will be a critical use of the optogenetic LOV2–BAX2 system in future studies.

## Materials and Methods

### Cell lines

HeLa cells were obtained from RIKEN BRC (RCB0007) and cultured in Dulbecco’s modified Eagle’s medium (DMEM) (D6546; Sigma-Aldrich) supplemented with 10% fetal bovine serum (FBS) (S1820-500; Biowest), 50 U/ml penicillin, and 50 µg/ml streptomycin (15070-063; GIBCO), at 37℃ in a 5% CO_2_ incubator.

### Plasmids

Plasmids for transient expression and stable expression in HeLa cells were generated by the Gibson Assembly method as follows. Gene sequences encoding EGFP, LOV2 (34), BAX (17), and the C-terminal regions of rat OMP25, human TMEM106B, and rat Cytb5 were inserted into the retroviral plasmid pMRX-IP (35). Halo, PLAAT3(C113S), Gal3, and FKBP, were inserted into the retroviral plasmid pMRX-IB (36). FKBP, FRB, mCherry-BAX(S184E) (17), TOM20, and LAMP1 were inserted into phCMV plasmid (17). EGFP, BAX, and the C-terminal regions of OMP25, TMEM106B, and Cytb5 were inserted into the lentiviral plasmid pLVX-TetOne-Puro (631849; Clontech) for doxycycline-dependent expression. The amino acid sequences of constructs used in this study are shown in Table S1.

### Transient expression by lipofection

HeLa cells were transiently transfected with the phCMV plasmid or pMRX-IP plasmid, both of which can be used for transient expression using Lipofectamine 2000 (11668019; Thermo Fisher Scientific).

### Stable expression by retrovirus or lentivirus infection

For retrovirus preparation, human embryonic kidney 293T (HEK293T) cells were transiently transfected with a retroviral plasmid, pCG-gag-pol, and pCG-VSV-G, (a gift from Dr. T. Yasui, National Institutes of Biomedical Innovation, Health and Nutrition, Osaka, Japan) using Lipofectamine 2000. For lentivirus, HEK293T cells were transiently transfected with a lentiviral plasmid, pCMV-VSV-G (gift from R.A. Weinberg, Whitehead Institute for Biomedical Research, Cambridge, MA), and psPAX2 (gift from D. Trono, Ecole Polytechnique Federale de Lausanne, Lausanne, Switzerland) using Lipofectamine 2000. After cells had been cultured for 3 days, the culture medium containing the virus was collected and passed through a 0.45-µm syringe filter unit. HeLa cells were incubated with the virus for 2 days in DMEM, and stable transformants were selected with 10 µg/ml puromycin (P8833; Sigma-Aldrich), 10 µg/ml blasticidin S (022-18713; Wako Pure Chemical Industries), or 200 µg/ml zeocin (R25001; Thermo Fisher Scientific).

### Doxycycline treatment

To enable doxycycline-dependent BAX mutant expression, HeLa cells were treated with 10 µg/ml doxycycline (D3447; Sigma-Aldrich) with 20 µM quinoline-Val-Asp-difluorophenoxymethylketone (Q-VD-Oph) (S1002; Selleck) and 200 nM Halo SaraFluor 650T Ligand (A308-01; Goryo Chemical) for 24–48 h.

### Heterodimerizer treatment

To induce heterodimerization of the FKBP and FRB tags, HeLa cells were treated with 500 nM A/C heterodimerizer (635057; Clontech).

### Blue light irradiation

HeLa cells were cultured in four-compartment glass bottom dishes (627870; Greiner Bio-One). Q-VD-Oph (S1002; Selleck) at 20 µM and Halo SaraFluor 650T Ligand (A308-01; Goryo Chemical) at 200 nM were added to DMEM 10 min before blue light irradiation. Blue light was irradiated from the bottom of the dish for 60 min using an LED plate (LEDA-B LED array and LAD-1 LED array driver; Amuza) at 37℃, with the following parameters: wavelength of 470 nm, a distance of 5 mm between the LED plate and the dish, a voltage of 13.5 V, an estimated luminosity level of 13.5 mW/cm^2^, in constant mode. Following irradiation, HeLa cells were incubated for 30 min in the dark, and then fixed with 4% paraformaldehyde (PFA) in phosphate-buffered saline (PBS) for 30 min. The PFA was removed, and the cells were washed three times with PBS.

### Fluorescence microscopy

Fluorescence microscopy images were acquired using a confocal laser microscope (FV3000; Olympus) equipped with a 60× oil-immersion objective lens (NA = 1.4) (PLAPON60XOSC2; Olympus). Images were captured using FluoView (Olympus). The software tool ImageJ was used for the analysis of the captured images.

### Correlative electron and light microscopy

For observation of mitochondrial morphology, HeLa cells expressing EGFP–LOV2– BAX(mito) and Halo–PLAAT3(C113S) were cultured on gridded coverslip-bottom dishes (TCI-3922-035R-1CS, a custom-made product based on 3922-035, with a cover glass attached in the opposite direction; Iwaki) coated with carbon by a vacuum evaporator (IB-29510VET, JEOL). The cells were stimulated by blue light for 60 min and incubated for an additional 30 min, followed by fixation with freshly prepared 2% PFA with 0.5% glutaraldehyde in 0.1 M phosphate buffer (pH 7.4). After washing with 0.1 M phosphate buffer three times, cells were observed with the FV3000 confocal laser scanning microscope system (Olympus), and then postfixed overnight with 2.5% glutaraldehyde in 0.1 M sodium cacodylate buffer at 4°C. After washing with 0.1 M sodium cacodylate, cells were treated with ferrocyanide-reduced osmium tetroxide (1% (*w*/*v*) OsO_4_, 1.5% (*w*/*v*) K4[Fe(CN)_6_]) in 0.065 M sodium cacodylate buffer for 2 h at 4°C, and rinsed five times using Milli-Q water. The samples were then stained with 2% uranyl acetate solution for 1 h and rinsed five times using Milli-Q water. The samples were dehydrated with an ethanol gradient series, covered with an EPON812-filled plastic capsule, which was placed inverted over the sample surface, and polymerized at 40°C for 12 h and then 60°C for 48 h. After polymerization, cover glasses were removed by soaking in liquid nitrogen, and the sample block was trimmed to about 150 × 150 μm, retaining the same area from which the fluorescence microscopy image was obtained. Then, serial sections (25 nm thick) were cut, collected on a silicon wafer strip, and observed under a scanning electron microscope (SEM; JEM7900F; JEOL).

### Quantification and statistical analysis

To calculate the percentage of mitochondria-ruptured cells and ER-ruptured cells, cells with colocalization of organelle markers and membrane-damage markers were counted manually by a researcher (who was blind to both the constructs and photostimulation of each sampled). For quantification of the number of Halo-Gal3 puncta per cell, the brightness of the Halo signal in each cell was adjusted by the ImageJ brightness filter and then binarized with automatic parameters. Puncta (size = 0.05–0.50 μm^2^, circularity = 0.10–1.00) were extracted using the “Analyze Particles” filter in ImageJ. Differences were statistically analyzed by one-way ANOVA. Statistical analysis was performed using Graph-Pad Prism 8 software.

## Acknowledgments

The authors thank Mr. Shoji Yamaoka for providing pMRX-IP; Mr. Teruhito Yasui for providing pCG-VSV-G and pCG-gap-pol; Ms. Yoko Ishida and Ms. Keiko Igarashi for their technical support for EM observation; and the Mizushima lab members for helpful suggestions and discussions.

## Data availability

All data supporting the analyses in the manuscript are available from the corresponding author upon reasonable request.

## Author contributions

Conceptualization: T.E., N.M.; Methodology: I.K.H.; Formal analysis: Y.N.; Investigation: Y.N.; Data curation: Y.N.; Writing original draft: Y.N., T.E., N.M.; Supervision: T.E., N.M.; Funding acquisition: T.E., N.M.

---

**Table S1.**
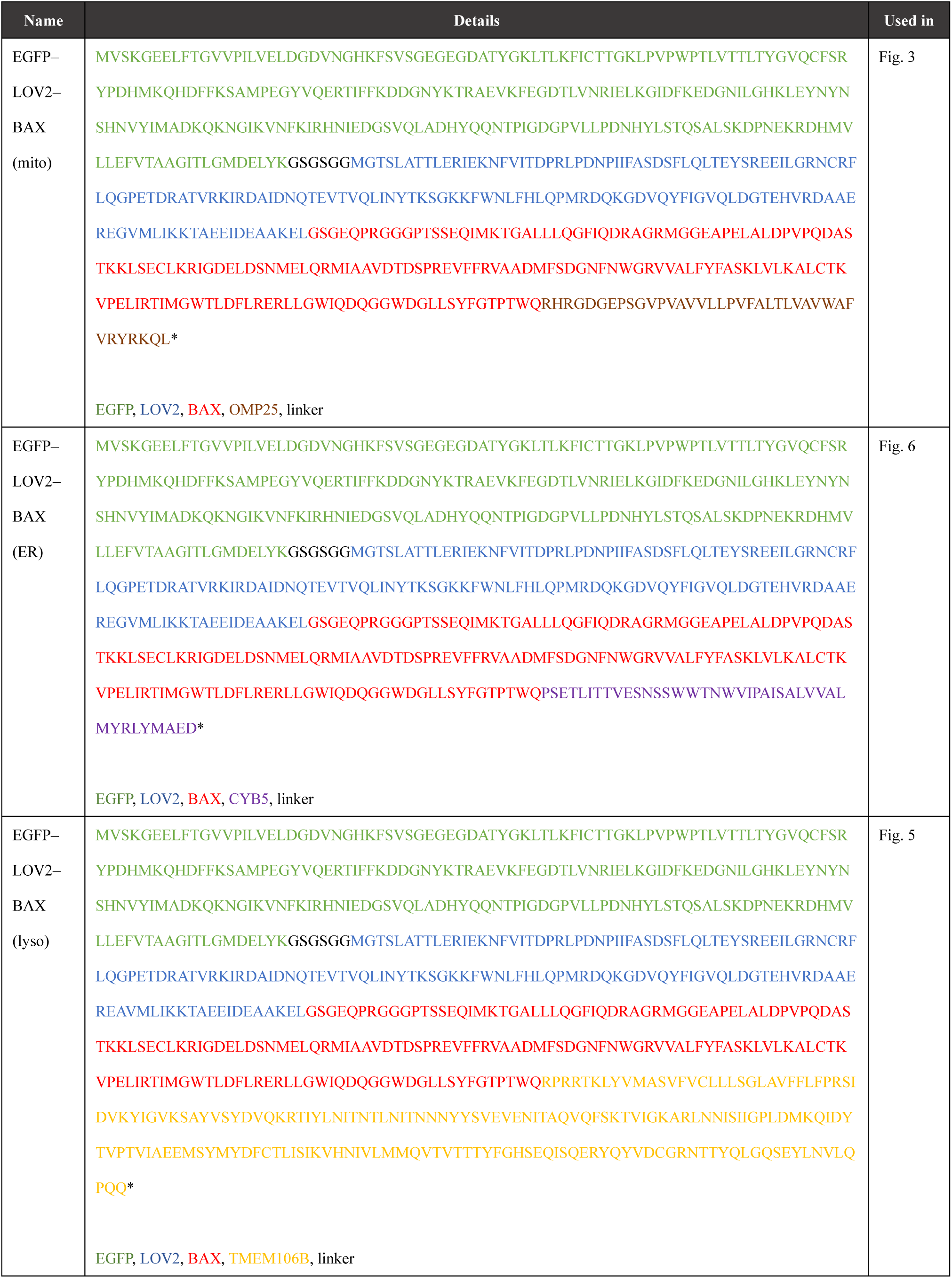
The amino acid sequences of constructs used in this study.

